# Magnetically Labeled iPSC-Derived Extracellular Vesicles Enable MRI/MPI-Guided Regenerative Therapy for Myocardial Infarction

**DOI:** 10.1101/2025.03.02.641040

**Authors:** Wenshen Wang, Zheng Han, Safiya Aafreen, Cristina Zivko, Olesia Gololobova, Zhiliang Wei, Geoffrey Cotin, Delphine Felder-Flesc, Vasiliki Mahairaki, Kenneth W. Witwer, Jeff W.M. Bulte, Robert G. Weiss, Guanshu Liu

## Abstract

Stem cell-derived extracellular vesicles (EVs) offer a promising cell-free approach for cardiovascular regenerative medicine. In this study, we developed magnetically labeled induced pluripotent stem cell-derived EVs (magneto-iPSC-EVs) encapsulated with superparamagnetic iron oxide (SPIO) nanoparticles for image-guided regenerative treatment of myocardial infarction, in which EVs that can be detected by both magnetic resonance imaging (MRI) and magnetic particle imaging (MPI). iPSC-EVs were isolated, characterized per MISEV2023 guidelines, and loaded with SuperSPIO20 nanoparticles using optimized electroporation conditions (300 V, 2 × 10 ms pulses), achieving a high loading efficiency of 1.77 ng Fe/10^6^ EVs. In vitro results show that magneto-iPSC-EVs can be sensitively detected by MPI and MRI, with a detectability of approximately 10^7^ EVs. In a mouse myocardial ischemia-reperfusion model, intramyocardially injected magneto-iPSC-EVs (2 × 10^9^) were imaged non-invasively by in vivo MPI for 7 days and ex vivo MRI, with the presence of magneto-iPSC-EVs confirmed by Prussian blue staining. Therapeutically, both native and magneto-iPSC-EVs significantly improved cardiac function, with a 37.3% increase in left ventricular ejection fraction and 61.0% reduction in scar size. This study highlights the potential of magneto-iPSC-EVs as a cell-free approach for cardiovascular regenerative medicine, offering both non-invasive imaging capabilities and therapeutic benefits for myocardial repair.

## Introduction

Cardiovascular disease (CVD) remains the leading cause of death and a global public health burden, with more than 1.5 million myocardial infarctions (MI) in the US annually. MI occurs when the cardiac muscle suffers prolonged ischemic injury following coronary occlusion. Even when coronary blood flow is restored, a subsequent inflammatory response often arises and inflicts additional damage, known as myocardial ischemia-reperfusion injury^1,2^. Current treatment strategies following MI are for early restoration of blood flow and to limit the ensuing adverse injury, chamber dilatation and impaired contractile function. Effective treatment of severely injured myocardium is still elusive^3^, in part because the adult heart has a limited number of stem cells and hence limited capacity of self-repair and regeneration.

Stem cell-based therapies have been extensively studied to treat heart diseases in the last two decades^4,5^, and a few clinical trials have been reported^6,7^. Mounting evidence now suggests that stem cells repair the injured myocardial cells predominately through a paracrine effect mediated by extracellular vesicles (EV), and hence stem cell-derived EVs may be used alone for cell-free regenerative medicine^8,9^. EVs are small membranous blebs or vesicles released from nearly all cell types and function as important messengers and mediators for intercellular communication in a diverse range of biological processes^10,11^. Depending on their biogenesis^12^, EVs are typically classified into exosomes (50–150 nm in diameter), microvesicles (50-500 nm), and apoptotic bodies (1–5 µm). Exosomes are derived from specialized intracellular compartments, *i.e.*, endosomes or multi-vesicular bodies (MVBs), microvesicles are shed directly from the plasma membrane, whereas apoptotic bodies are released by dying cells. When reaching recipient cells that are either in the immediate vicinity or at a distance, EVs fuse with them to transmit cargo (*e.g.,* proteins, messenger RNA, microRNA, and lipids) and elicit biologic changes in recipient cells^13^. EVs secreted by stem cells have been tested for regenerative potential in different disease models, including myocardial injury^14,15^. While the mechanisms are incompletely understood, it is believed that EVs exert protective and/or reparative effects through cytoprotection, stimulation of angiogenesis, induction of antifibrotic cardiac fibroblasts, and modulation of M1/M2 polarization of macrophages infiltrating the infarcted region^16,17^. Compared to their parental stem cells, human stem cell-derived EVs are considered to be a safer and more effective regenerative medicine approach for treating otherwise untreatable cardiac disease^14,18,19^.

Besides encouraging preclinical studies, rapid progress has been made in translating EVs to human studies. Currently, encouraging clinical results from a Phase II trial (NCT04493242) were reported^20^. In that study, an investigational new drug (IND), ExoFlo™ (EVs derived from bone marrow-derived mesenchymal stem cells, MSC-EVs), exhibited favorable safety data and significant efficacy in treating hospitalized adult COVID-19 patients with moderate-to-severe acute respiratory distress syndrome^20^. In addition, there are over 60 interventional studies utilizing EVs as therapeutic agents, with 40 of them involving stem cell-derived EVs^21,22^, including two phase III trials (NCT05813379 and NCT05969717)^20^.

In vivo tracking of EVs is indispensable for developing, testing and optimizing EV therapy by providing information about the location and quantity of the administered EVs. There have been significant efforts to develop EV tracking methods in the last decade^23^. Among them, optical imaging is the most widely used imaging modality, including fluorescence imaging^24,25^ and bioluminescence imaging^26^, owing to its inherent high sensitivity and wide availability in preclinical labs. However, the application of optical imaging to deep organs in large animal models and in humans is often challenging. Other imaging modalities such as single-photon emission computed tomography (SPECT)^27^, positron emission tomography (PET)^28^, computed tomography (CT)^29,30^, magnetic resonance imaging (MRI)^31,32^, and magnetic particle imaging (MPI)^33,34^ have also been reported. Among them, MRI is an appealing imaging modality widely available in the clinic, has excellent soft tissue contrast and no ionizing radiation. MRI can provide high spatial resolution images of the location and relative quantities of EVs^31,32^ by labeling them with superparamagnetic iron oxide particles (SPIOs). Besides MRI, loading cells or EVs with SPIOs also makes it possible to detect them using magnetic particle imaging (MPI)^35^, an emerging tracer-based imaging technology that has ultra-high sensitivity and hot-spot (zero background signal) specificity^36–38^. We have recently developed a novel strategy to prepare highly purified, magnetically labeled EVs, termed magneto-EVs, using surface-modified, “sticky” SPIO, offering an efficient purification procedure that is less equipment-intensive, and much faster (∼min), to prepare EVs with excellent purity and high SPIO loading. With this technical advance, we demonstrated the first MRI tracking of the delivery of therapeutic EVs to the sites of injury using a systemic injection route^39^.

We aimed to establish an innovative theranostic platform that combines advanced imaging capabilities with therapeutic stem cell-derived EVs for precision treatment of MI. Building upon our previous development of magneto-EVs, a novel magnetic labeling approach that enables MRI tracking of iPSC-derived EVs to injury sites^39^, we now advance this technology in two significant directions. First, we optimized the labeling protocol to achieve dual-modality imaging with both MPI and MRI, enhancing our ability to quantitatively assess EV biodistribution with greater precision. Second, we rigorously evaluate the potency and therapeutic efficacy of these magneto-EVs through a comprehensive panel of in vitro and in vivo assays, confirming their regenerative properties. This integrated approach represents a crucial step toward realizing image-guided, EV-based regenerative therapies for cardiac repair.

## Methods

### Materials

Dendronized superparamagnetic iron oxide (SPIO) nanoparticles (core diameter = 20 nm, SUPERSPIO20@D1-2P) were obtained from SuperBranche (Strasbourg, France). Hexa-histidine peptide was purchased from GenScript (Piscataway, NJ, USA). All other chemicals, including 1-Ethyl-3-(3-dimethylaminopropyl)-carbodiimide hydrochloride (EDC, cat# E6383), N-hydroxysulfosuccinimide sodium salt (sulfo-NHS), and Ni-NTA were purchased from Sigma-Aldrich (St. Louis, MO, USA). Size exclusion chromatography was performed using qEV1 columns (70 nm) purchased from iZON Science (Cambridge, MA, USA). Protein quantification was carried out using the Pierce™ Micro BCA™ Protein Assay Kit (cat# 23235) from Thermo Scientific (Waltham, MA, USA). RNA quantification was performed using the Quant-iT™ RiboGreen™ RNA Assay Kit (cat# R11490) from Invitrogen (Thermo Fisher Scientific, Eugene, OR, USA).

### Cells

Human BC1-induced pluripotent stem cells (iPSCs), originally reprogrammed from CD34+ hematopoietic progenitor cells of a healthy male donor ^40^ were maintained in Essential 8™ (E8) medium (Gibco, cat# A1517001) supplemented with 10 µM Y-27632 ROCK inhibitor (Ri, STEMCELL Technologies, cat# 72304)^41,42^. Between 3^rd^ −12^th^ passages after thawing, cell culture media (CCM) was collected on days 2 and 3 for EV isolation. Representative microscopy images of the BC1-iPSCs are shown in **Figure S1**.

### EV isolation and characterization

EVs were isolated using the method previously described^39^. Briefly, human iPSC-conditioned culture medium (CCM) was centrifuged at 300 × g for 10 minutes, followed by a second centrifugation at 2,000 × g for an additional 10 minutes at 4 °C to remove cells and debris. The supernatant was then concentrated using an Amicon Ultra-15 centrifugal filter (100 kDa) column by centrifugation at 4,000 × g for 20 minutes (MilliporeSigma, Billerica, MA, USA). Each time, 15 mL of medium was loaded onto the column, yielding approximately 400 μL of retentate. The concentrated EV solution was subsequently purified by size exclusion chromatography (SEC) using qEV1 (70 nm) columns (iZON, Cambridge, MA, USA). After washing the qEV1 columns with 10 mM PBS, pH=7.2-7.4, 0.5-1.0 mL of the concentrated EVs was applied to the top of the column and eluted with PBS. A total of 2.8 mL of EV-rich fractions (4 × 0.7 mL) were collected. The purified EVs were further concentrated using an Amicon column, and the final volume was adjusted to achieve a concentration of approximately 1 × 10^11^ EVs per mL. The number of EVs was quantified using Nano Flow Cytometry (NanoFCM Inc.) with NanoFCM software (Profession V1.0). The isolated EVs were characterized according to MISEV2023^43^ with respect to both EV biomarkers (CD9) and non-EV biomarkers (Calnexin) (**Figure S2)**. The size of EVs was measured by DLS (Nanosizer ZS90, Malvern Instruments) and transmission electron microscopy (TEM) using a Zeiss Libra 120 TEM operated at 120 KV and equipped with an Olympus Veleta camera (Olympus Soft Imaging Solutions GmbH, Münster, Germany).

### Magnetic labeling of EVs

<u>SuperSPIO20-His</u> was synthesized using a modified protocol adapted from our previous method^39^. In brief, 400 μL of dendronized SPIO (SuperSPIO20, 1 mg/mL,) was mixed with 40 μL of EDC/NHS solution (1 mg/mL) and 560 μL of MES buffer (pH=6.0). The mixture was incubated at room temperature (RT) with gentle shaking for 30 minutes to activate the carboxyl groups. Following activation, 1 mg of hexa-histidine peptide (dissolved in 100 μL PBS) was added to the reaction mixture, and the pH was adjusted to 7.4, followed by incubation at RT with gentle shaking for 2 hours. The resulting conjugate was purified using an Amicon Ultra-100K column by washing three times with ddH2O.

<u>Electroporation</u> was performed using a Celetrix EX+ electroporator (Celetrix, Manassas, VA) using a modified procedure as described previously^39^. In brief, 40 μL of iPSC-EVs (2 × 10^10^ EV/mL) and 10 μL of SuperSPIO20-His (2 mg/mL) were mixed with 50 μL of buffer of choice and loaded into a vertical electroporation cell (100 µL, path length=10 mm) for electroporation. The sample was subsequently purified using an Ni-NTA column to remove unencapsulated SPIOs. The iron content in the purified EV product was determined by measuring UV absorption at 500 nm^32^.

<u>The protective effect</u> of different electroporation buffers on electroporation-induced damage of EVs was first investigated. Tested buffers were a) 10 mM potassium-phosphate buffer (KPBS, pH=7.4), b) KPBS containing 50 mM sucrose, c) KPBS containing 50 mM trehalose^44^, and d) Celetrix electroporation buffer supplied by the electroporator manufacturer^45^.

Based on our initial testing, KPBS solution containing 50 mM trehalose was chosen in all the following studies. To optimize the electroporation conditions, the loading rate and impact of EV size by the voltage, length, and number of the applied pulses were investigated in the following combinations: a) 200V/250V/300V/350V/400V with 10 ms and 1 pulse, b) 2 ms/5 ms/10 ms/20 ms with 300V and 1 pulse, c) 1 pulse/2 pulses/5 pulses/10 pulses with 300V and 10 ms. The vesicular protein and RNA were assessed before and after electroporation using a MicroBCA protein assay kit and a RediPlate™ 96 RiboGreen™ RNA quantitation kit after centrifugation with a 100K Amicon Ultra-15 filter, according to the manufacturer’s instructions.

The SuperSPIO20-His bound to the Ni-NTA column was eluted using a buffer composed of 50 mM NaH_2_PO_4_, 300 mM NaCl, and 2.5 M imidazole (pH = 9.2). To ensure efficient dissociation of the SuperSPIO20-His from the column, the elution solution was incubated at 40°C for 1 hour before collecting the elute of recovered SuperSPIO20-His.

### Stability of vesicular SuperSPIO20

To evaluate the stability of SuperSPIO20 encapsulated within EVs, the retention of SuperSPIO20 in magneto-EVs was assessed at various time points over a 7-day period. In brief, magneto-EVs were incubated in PBS at room temperature or in fetal bovine serum (FBS) at 37°C at a volume ratio of 1:2 (total volume = 100 μL). At each time point, free SuperSPIO20-His released from the vesicles was removed using Ni-NTA purification, and the remaining vesicular SuperSPIO20 content was quantified using a Ferrozine-based colorimetric assay^46,47^.

### RNA-Seq library preparation and sequencing

Total RNA from native iPSC-EVs and magneto-iPSC-EVs was isolated using TRIzol (Sigma) and the RNeasy® Mini Kit (Qiagen) following the manufacturer’s protocols. Exosomal small RNA next-generation sequencing (NGS) was conducted by Novogene Corporation Inc. (Sacramento, CA, USA). Small RNA tags were aligned to the reference genome using Bowtie with no mismatches allowed to analyze their expression and distribution. Known miRNAs were identified by mapping the aligned tags to miRBase 20.0 using modified software tools, including mirdeep2 and srna-tools-cli^48^. Differential expression analysis between native iPSC-EVs and magneto-iPSC-EVs was performed using the DESeq R package (v1.8.3). P-values were adjusted for multiple comparisons using the Benjamini & Hochberg method, with an adjusted P-value threshold of 0.05 set to determine significant differential expression.

### Hypoxia/reoxygenation (H/R) assay

Rat cardiac H9c2 cells were used as a cellular H/R model to evaluate the cardiac protective effects of EVs. H9c2 cells (American Type Culture Collection) were cultured in DMEM containing high glucose supplemented with 10% FBS (Thermo Fishier Scientific, Inc.) and 100 U/mL streptomycin-penicillin at 37°C in a 5% CO_2_ atmosphere. The cellular H/R model was established by culturing H9c2 cells in DMEM containing low glucose without FBS and with 5% CO_2_ in a 1% O_2_ (hypoxic) atmosphere for 6h, followed by culturing in normal 20% oxygen for 48 h in the presence of native or magneto-iPSC-EVs (1×10^9^ per mL). The ratio of apoptotic cells was measured using flow cytometry after cells were stained with an Annexin V/PI kit according to the manufacturer’s instructions.

### qRT-PCR

To evaluate the anti-inflammation effects of EVs, their effects on M1-polarized macrophages were assessed. In brief, 1 × 10^5^ murine RAW 264.7 macrophages (ATCC #TIB-71) were seeded into 12-well plates and cultured for 3 days. M1 polarization was induced by incubating cells with lipopolysaccharide (LPS; 100 ng/mL, Sigma) and interferon-gamma (IFN-γ; 10 ng/mL, R&D Systems) for 24 hours, according to Boehler et al^49^. After polarization, the cells were washed and switched to fresh media containing PBS, 10^7^ native iPSC-EVs, or 10^7^ magneto-iPSC-EVs, respectively, and incubated for an additional 48 hours. RAW 264.7 cells were collected and homogenized in 500 µL of TRIzol reagent (Sigma) to extract total RNA using the RNeasy® Mini Kit (Qiagen) according to the manufacturer’s instructions. RNA concentration was measured using a Nanodrop 2000 spectrophotometer (Thermo Fisher Scientific). Complementary DNA (cDNA) was synthesized from the extracted RNA using the SuperScript III First-Strand Synthesis System (Invitrogen). The cDNA served as a template for gene amplification via qRT-PCR using the CFX96™ Real-Time PCR System (Bio-Rad). Primer sequences used for qRT-PCR are listed in **Table S1**. The PCR cycling conditions consisted of 40 cycles at 95 °C for 3 seconds, 60 °C for 15 seconds, and 72 °C for 15 seconds. To quantify RNA expression and assess cDNA synthesis efficiency, target gene levels were normalized to the expression of the endogenous reference gene 18S ribosomal RNA (18S rRNA).

### *In vitro* MRI and MPI

The T_2_ MR relaxation time of magneto-EVs at different concentrations (0.5 – 8×10^9^ EV/mL) in PBS was measured using a Carr-Purcell-Meiboom-Gill (CPMG) method at room temperature on an 11.7T Biospec (Bruker) vertical bore MRI scanner according to our previously published procedure^50^. The acquisition parameters were TR/TE= 25,000/4.3 ms, RARE factor =16, matrix size = 64 × 64, in-plane resolution =0.25 × 0.25 mm, and slice thickness = 2 mm. All data were processed using Matlab. The T_2_ relaxivities (r_2_) were calculated using the following equation: R_2_ = R_20_+ r_2_ × C, where R_20_ represents the inherent water transverse relaxation rate (R_2_=1/T_2_).

Momentum™ MPI scanner (Magnetic Insight Inc., Alameda, CA, USA) was used to characterize the MPI properties of SuperSPIO20 and magneto-iPSC-EVs. The sensitivity and resolution were characterized using the RELAX module. Particle sensitivity was defined as the peak amplitude of the point spread function (PSF) normalized to iron content (μg). The resolution was measured as the full-width half-maximum (FWHM) of the PSF. 2D coronal MP images were also acquired and analyzed using ImageJ/Fiji (NIH, Bethesda, MD).

### Animal model and treatment

All animal experiments were conducted in compliance with protocols approved by our Institutional Animal Care and Use Committee. Female C57BL/6J mice (6–8 weeks old) were obtained from Jackson Laboratories (Bar Harbor, ME, USA). Cardiac ischemia-reperfusion injury (IRI) was induced by a 35-minute ligation of the left anterior descending (LAD) coronary artery, followed by reperfusion according to previously described^51^, which is detailed in the **Supporting Information**. Immediately after reperfusion, treatment was administered via a three-site intramyocardial injection at a total dose of 2 × 10^9^ EVs per mouse. Mice were randomly assigned to receive saline, iPSC-EVs, or magneto-iPSC-EVs, with a final sample size of 10-12 per group. The variation in group size was due to some mice not surviving the procedure. A sham control group (n=4) was also included for comparison.

### In vivo MPI

All in vivo MP imaging was performed using a Momentum™ MPI scanner. At 1 and 7 days after magneto-iPSC-EV administration, mice were scanned in 3D imaging mode using the *high sensitivity* configuration. The imaging parameters included an excitation field amplitude of 5 mT, a gradient field strength of 5.7 T/m, and a drive frequency of 45 kHz. Each 3D scan consisted of 21 projections (total acquisition time = 20 min). To facilitate image alignment, two fiducial markers (2 µg/mL SuperSPIO20 solution, 100 µL in microcentrifuge tubes) were placed on the sample holder alongside the mouse.

After each MPI scan, the mouse was transferred to a micro-CT scanner while being secured on the sample holder using a custom-designed 3D-printed adapter. CT scans were acquired using an IVIS Spectrum CT In Vivo Imaging System (Perkin-Elmer, Shelton, CT, USA), with 720 projections and 20 ms exposure time each. The scanning area was adjusted to ensure that both the mouse and fiducial markers were included within the field of view. The use of the same holder for both MPI and CT imaging allowed co-registration of MPI and CT images, ensuring precise spatial alignment for accurate anatomical and functional analysis. After the final MPI scan, mice were sacrificed to harvest the heart. After rinsing with PBS, the excised hearts were fixed in 4% paraformaldehyde (PFA) at 4 °C for at least 24 h.

### Ex vivo MRI

Each fixed heart was transferred to a 15 mL tube filled with hydrogen-free Fomblin (Solvay Solexis, Inc., USA) and assessed for ex vivo high-resolution MRI using a vertical bore 11.7T Bruker scanner equipped with a 20 mm birdcage transmit/receive volume coil. Imaging was performed with a three-dimensional FLASH sequence, using parameters of TE = 6 ms, TR = 150 ms, matrix size = 310 × 240 × 145, field of view (FOV) = 12.06 × 9.48 × 5.68 mm, resolution = 39 × 39 × 39 μm³, 5 averages, and a flip angle of 15°. MR images were first processed using ImageJ, and 3D visualization of the distribution of magneto-EVs was performed using 3D Slicer Software 5.2.2.

### Cardiac MRI

On day 30 following treatment, each mouse underwent a multi-slice cine cardiac MRI using a self-gated FLASH sequence (IntraGate FLASH, ParaVision 6.0). For data analysis, left ventricular volumes were assessed by manually delineating the endocardial and epicardial boundaries on short-axis slices using AnimHeart software (CASIS, Dijon, France). This delineation was performed on both end-diastolic and end-systolic frames to calculate the end-diastolic volume (EDV) and end-systolic volume (ESV). To quantitatively evaluate cardiac function, the left ventricular ejection fraction (LVEF) was calculated using the formula: (EDV-ESV)/ESV × 100%.

### Histopathology

#### Prussian blue staining

Mouse hearts were paraffin-embedded and sectioned into 25 µm thickness slices after *ex vivo* MRI. Prussian blue staining was performed according to a previously published protocol^39^ to detect iron deposits within the tissue^39^.

#### Masson’s Trichrome staining

After the cardiac MRI evaluation, mouse hearts were harvested, fixed in paraformaldehyde fixative solution (Thermo Scientific), and embedded in OCT compound and snap-frozen at −80°C. Using a cryostat (CryoStar™ NX70, Thermo Fisher Scientific, UK), 25 µm thick sections were cut, starting 250 µm from the apex. The tissue sections were stained with a Masson’s Trichrome kit (Abcam, ab150686) according to the manufacturer’s protocol. All microscopic imaging was conducted using a Leica Thunder imaging system equipped with a DFC-9000-GT camera (Leica Microsystems).

Fibrosis area was quantified from Masson’s Trichrome-stained sections using ImageJ software (NIH). The fibrotic (blue) and viable myocardium (red) areas were segmented and measured, and the fibrosis percentage was calculated.

### Statistical analysis

All data are presented as mean ± SD unless otherwise noted. Statistical analysis was performed using GraphPad Prism version 10 (GraphPad Software Inc., San Diego, CA, USA). The D’Agostino–Pearson test was conducted to assess the normal distribution of data for all groups except the sham control group (n=4), for which the Shapiro–Wilk test was used. An unpaired two-tailed Student’s t-test was utilized to compare two parametric groups. One way Analysis of Variance (ANOVA) was used to determine the differences between the three groups of mice receiving different treatments (vehicle control, iPSC-EVs, and magneto-iPSC-EVs). The Mann– Whitney U test was employed to compare the difference between a group and the sham group, which did not pass the normality test. A p-value of < 0.05 was considered statistically significant.

## Results

### iPSC-EV production

We have developed and optimized the procedures for iPSC-EV production as shown in **Figure 1**. We chose to use concentrated commercial size exclusion chromatography (SEC), *i.e.*, qEV columns by iZON for reproducible EV isolation. Three EV-rich fractions (7–9, 0.5 mL each) were pooled. The final concentration was ∼ 4×10^10^ EVs/mL, derived from 105 mL CCM (approximately 17-29 million cells). The protein and RNA contents were determined to be 19.7 µg and 400 ng per 10^10^ EVs with small batch-to-batch variations (coefficient of variation < 10%). The quality and purity of the EV products were confirmed by EV markers (transmembrane protein CD9) and non-EV markers (Calnexin) (**Figure S2**) according to MISEV2023 guidelines^43^.

**Figure 1.**
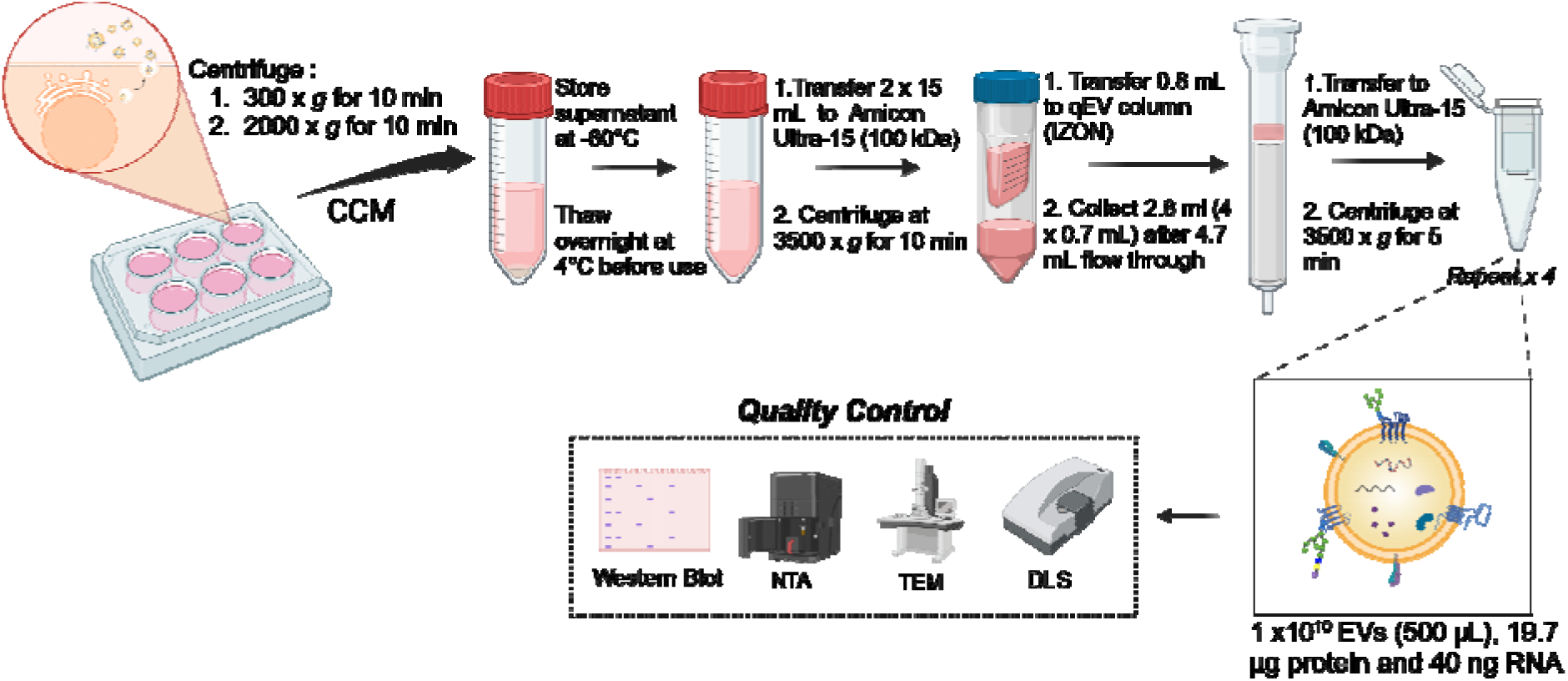
Schematic illustration of EV isolation and characterization.

### Optimizing magnetic labeling for sensitive MPI and MRI detection of iPSC-EVs

To produce magneto-EVs with high MRI and MPI detectability, we optimized the electroporation conditions to maximize SuperSPIO20 loading while minimizing the morphological and functional impacts. Before optimization, we conducted a comparative experiment to identify the buffer that best protects EVs during electroporation. Among the four buffers tested (**Figure S3a**), potassium phosphate buffer (KPBS) supplemented with 50 mM trehalose resulted in the smallest increase in EV size, indicating superior protective effects. Therefore, KPBS + 50 mM trehalose was selected for all subsequent experiments.

Next, we investigated the effects of voltage and pulse duration on SuperSPIO20 labeling efficiency and EV size. As shown in **Figures 2b,c**, both labeling efficiency and EV size increase approximately linearly with voltage. To balance these two factors, we calculated the ratio of labeling efficiency to EV size increase (**Figure 2d**), which initially increased and then decreased with higher voltages. Two “optimal values” were identified: one favoring higher loading efficiency and another minimizing size increase. Since maximizing MRI and MPI detectability was prioritized, we selected 300 V as the optimal voltage, providing high labeling efficiency while maintaining a relatively small size increase.

**Figure 2.**
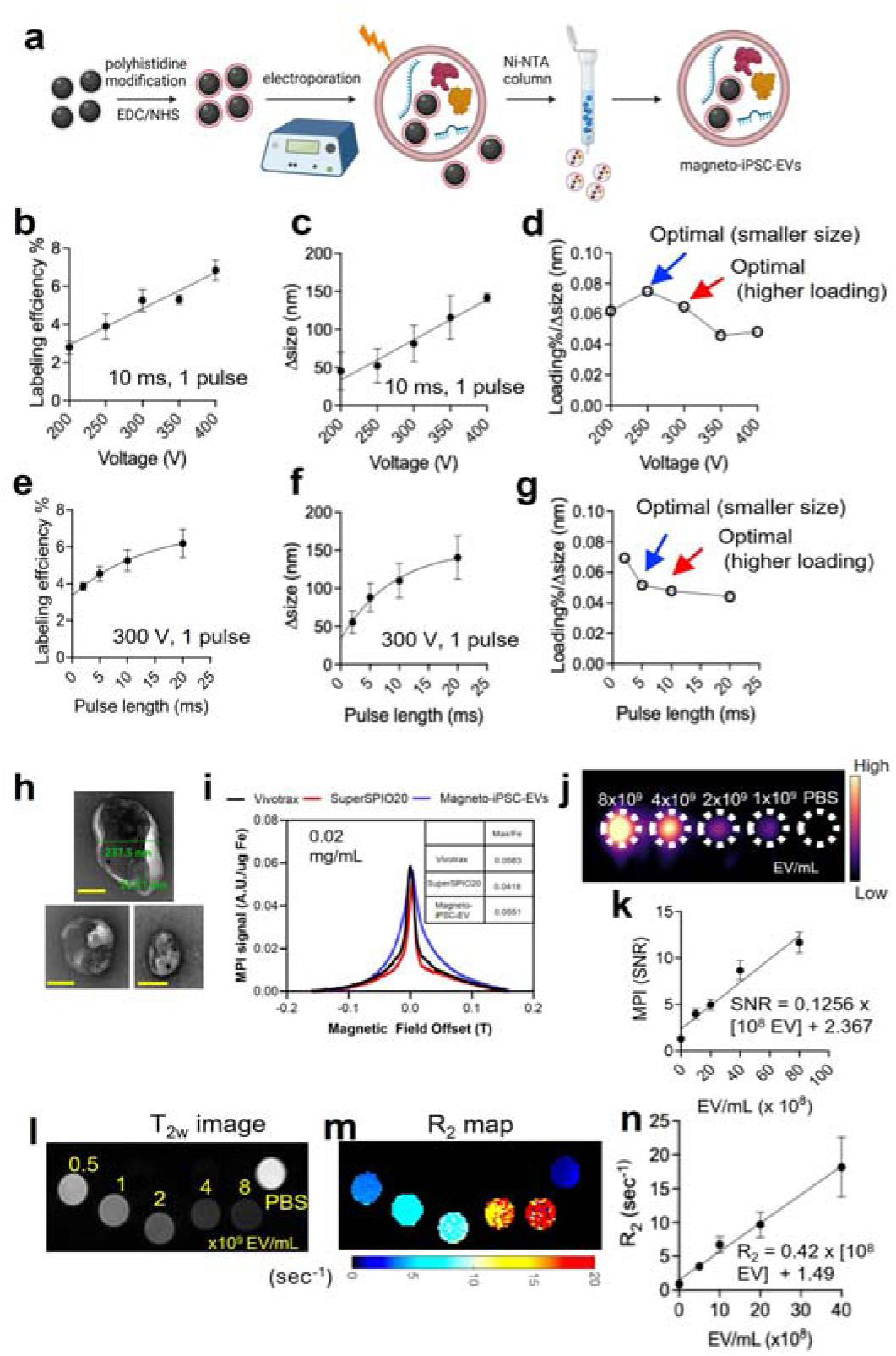
Preparation of magneto-EVs with enhanced MPI and MRI detectability. (**a**) Schematic illustration of magnetic labeling and subsequent purification of iPSC-EVs using surface-modified SuperSPIO20. In this study, we used hexa-histidine to coat SuperSPIO20, enabling efficient removal of free SuperSPIO20 in the solution via Ni-NTA columns, thereby producing magneto-iPSC-EVs with negligible contamination from unencapsulated SuperSPIO20 in the final product. (**b**) Labeling efficiency as a function of electroporation voltage (one pulse, length = 10 ms). (**c**) EV size increase as a function of electroporation voltage. (**d**) Determining the optimal voltage based on the ratio of labeling efficiency and size change. (**e**) Labeling efficiency as function of electroporation pulse length (one pulse, voltage = 300 V). (**f**) EV size increase as function of electroporation pulse length. (**g**) Determining the optimal pulse length based on the ratio of labeling efficiency and size change. (**h**) TEM images of magneto-iPSC-EVs (three different fields of view). Scale bar = 100 nm. (**i**) MPI relaxometry comparing the sensitivity and resolution of SuperSPIO20 (red) and SuperSPIO20-loaded EVs (blue), with Vivotrax (black) included as a reference. (**j**) MPI image of magneto-iPSC-EVs at varying concentrations. (**k**) Calibration curve of MPI signal versus the concentration of magneto-iPSC-EVs. (**l**) T2-weighted image and (**m**) R2 map of magneto-iPSC-EVs at different concentrations. (**n**) R2 relaxation rate as a function of the concentration of magneto-iPSC-EVs. All data are presented as mean ± SD from three independent experiments. Similarly, we analyzed the effect of pulse duration on labeling efficiency and EV size (**Figures 2e–g**). A pulse duration of 10 ms was determined to be optimal to provide high labeling efficiency. Interestingly, when varying the number of pulses while keeping voltage and total pulse duration the same (300 V and 10 ms), no significant difference in labeling efficiency was observed between 2 pulses and 10 pulses (**Figure S4**). Based on these findings, we established the optimal electroporation conditions for SuperSPIO20 loading to be 300 V, 10 ms, and 2 pulses. These parameters achieved high labeling efficiency without dramatically increasing EV size. Of note is that free SuperSPIO20-His particles removed during purification can be recovered and recycled for reuse. Using an elution buffer containing 50 mM NaH_2_PO_4_, 300 mM NaCl, and 2.5 M imidazole, approximately 25% of SuperSPIO20-His was successfully recovered from the Ni-NTA column by each elution. Hence, this recycling process can significantly reduce the overall cost of magneto-EV production.

Using the optimized electroporation conditions (300 V, 10 ms, 2 pulses), we achieved a SuperSPIO20 loading efficiency of 1.77 ng iron per 10^6^ iPSC-EVs, corresponding to a loading rate of 6.9%, with a moderate impact on particle size (a 46% size increase). Transmission electron microscopy (TEM) images (**Figure 2h**) confirmed the well-preserved morphology of magneto-EVs, with an average size of 237.5 nm (SuperSPIO20 core size = 25.3 nm). Notably, many EVs were observed to be loaded with multiple SuperSPIO20 particles. After electroporation, the protein content was measured to be 95.0± 6.4%, while the RNA content was 86.6± 12.1% (**Figure S5**), indicative of well-reserved EV functionalities.

The magnetic labels were revealed to be relatively stable. As shown in **Figure S6**, incubating in PBS at ambient temperature for 7 days resulted in approximately 15% of SuperSPIO20 released from magneto-iPSC-EVs, indicating a relatively stable shelf life. In a similar manner, 78% of SuperSPIO20 was measured to be retained in EVs after incubation in serum at 37 °C for 7 days, indicating a slow in vivo release rate.

An in vitro phantom MPI experiment was conducted to address two key questions: (1) the detectability of magneto-EVs using MPI and (2) whether MPI provides sufficient sensitivity for in vivo quantification. Relaxometry results showed that magneto-iPSC-EVs produced a high MPI signal (0.0551 AU/µg Fe) that is slightly higher than free SuperSPIO20 (0.0418 AU/µg Fe) and is close to the commercial MPI tracer Vivotrax (0.0583 AU/µg Fe). The full-width at half-maximum (FWHM) of magneto-iPSC-EVs was slightly larger than that of SuperSPIO20 (0.05 vs. 0.04) (**Figure 2i**). There was a good linear relation between MPI signal intensity and EV concentration (r=0.92, **Figure 2j**). Based on a standard curve (**Figure 2k**), the detectability threshold (SNR= 3) for MPI was determined to be 1 × 10^7^ EVs for a 10 µL sample, corresponding to 1× 10^9^ EVs/mL.

In vitro MRI measurements were performed using magneto-iPSC-EV samples at concentrations ranging from 0.5 to 8.0 × 10^9^ EVs/mL (volume = 20 µL) (**Figure 2i-n**). The results indicated that the MRI detectability was as low as 2.5 x 10^8^ EVs/mL (for generating an MRI signal change of ΔR2= 1 s^−1^), corresponding to 0.5 × 10^7^ EVs in a sample volume of 20 µL. These findings demonstrate that both MPI and MRI provide robust sensitivity for detecting and quantifying magneto-iPSC-EVs at therapeutic concentration levels.

### Magneto-iPSC-EVs show well-preserved function and therapeutic potency

To determine the transcriptome profile of magneto-iPSC-EVs and the effects of electroporation, we performed a small RNA-Seq analysis as depicted in **Figure 3a**. Many highly expressed small RNAs in iPSC-EVs (both native and magneto-) have the functions related to anti-apoptosis and anti-inflammation (**Table S2**). Particularly, three small RNAs (i.e., miR-30d-5p, miR-26a-5p, and miR-423-5p) have been reported previously to be beneficial for myocardial infarction^52–54^. Importantly, the volcano plot analysis revealed only 16 miRNAs are different between the two groups (**Figure 3b**). The heatmap and Pearson correlation results indicated no significant gene expression changes across native and magneto-iPSC-EVs (**Figure 3c, d**). These results show that the profile of small RNA in magneto-iPSC-EVs was not significantly altered by magnetic labeling, confirming that magneto-iPSC-EVs preserve their therapeutic function.

**Figure 3.**
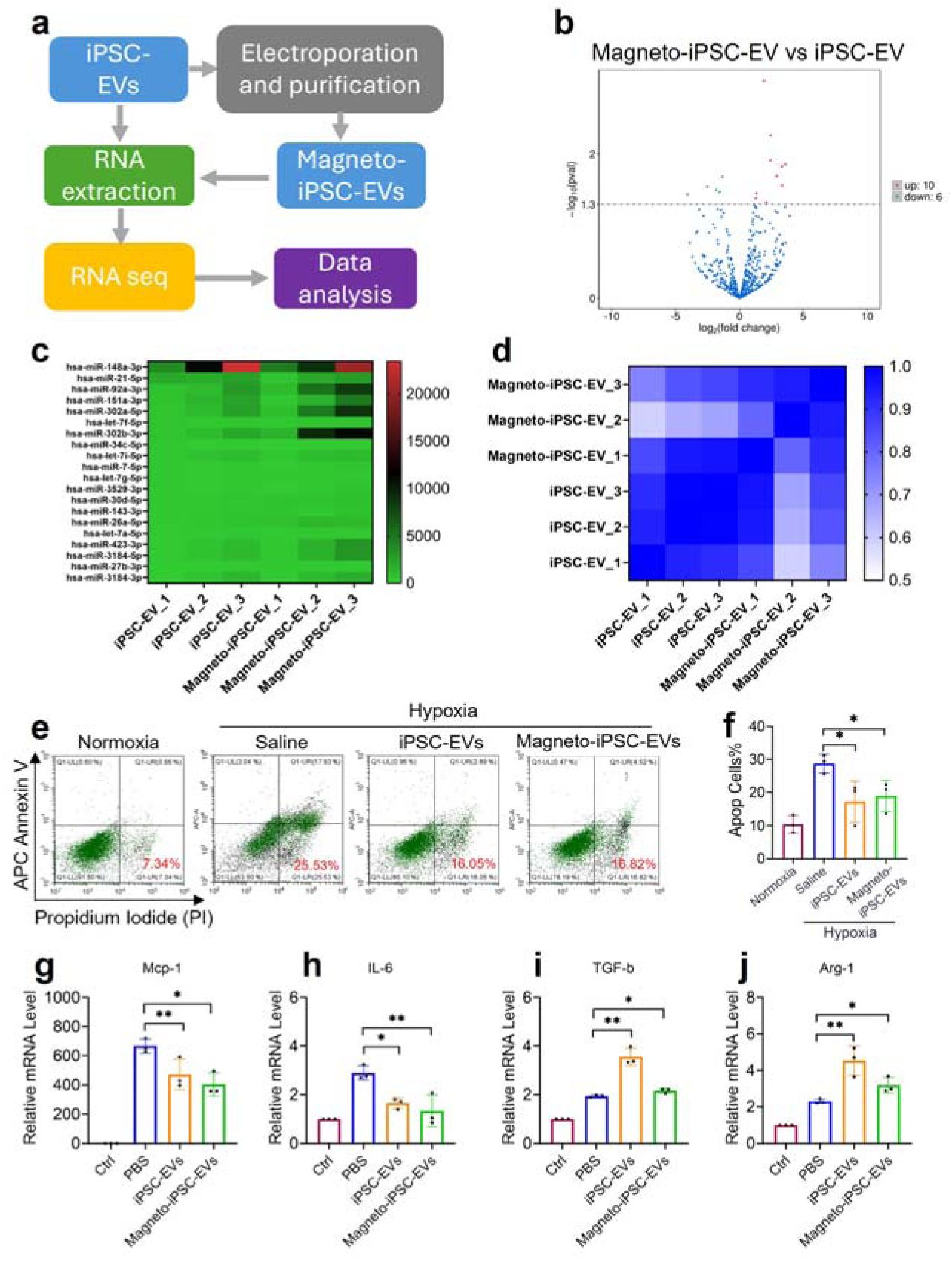
Transcriptome analysis and evaluation of in vitro therapeutic effects for native iPSC-EVs and magneto-iPSC-EVs. (**a**) Schematic workflow of RNA sequencing analysis comparing small RNA profiles between iPSC-EVs and magneto-iPSC-EVs. (**b**) Volcano plot depicting differential small RNA expression between magneto-iPSC-EVs and iPSC-EVs. (**c**) Heatmap of differentially expressed small RNAs between sample groups. (**d**) Correlation matrix showing the Pearson correlation coefficients between triplicates of iPSC-EV and magneto-iPSC-EV samples. (**e**) Representative flow cytometry analysis showing apoptosis patterns in H9c2 cardiomyocytes following hypoxia/reoxygenation and EV treatment. (**f**) Quantification of apoptotic cells demonstrating the protective effects of iPSC-EVs and magneto-iPSC-EVs against hypoxia-induced cell death (mean ± SD, n = 3, *P < 0.01). (**g-j**) Expression levels of inflammatory markers Mcp-1 (M1), IL-6 (M1), TGF-b (M2), and Arg-1 (M2) in LPS/ IFN-γ induced RAW264.7 macrophages following EV treatment as measured by qRT-PCR.

Next, we investigated the cardiac protective effects of magneto-iPSC-EVs on rat cardiac H9c2 cells under H/R conditions (**Figure 3e, f**). Cells treated with either native EVs or magneto-iPSC-EVs exhibited significantly lower apoptosis ratios compared to the control group. Specifically, the percentages of apoptotic cells were 19.0 ± 4.7% and 17.2 ± 6.3% in the magneto- and native iPSC-EVs treatment groups, respectively, while the control group showed 28.8 ± 2.9%. Although the native EVs demonstrated slightly higher protection than the magneto-EVs (**Figure 3f**), this difference was not significant. This experiment further confirmed the sustained cardiac protective effects of magneto-iPSC-EVs.

We also compared the immunomodulatory effects of native and magneto-iPSC-EVs on M1-polarized Raw264.7 macrophages stimulated by LPS/IFN-γ. The results show that magneto-iPSC-EV incubation resulted in upregulation of M2 markers (i.e., TGF-β, Arg-1, **Figure 3g, h**) and downregulation of M1 markers (i.e., Mcp-1, IL-6, **Figure 3i, j**), indicating that magneto-iPSC-EVs possess a similar immunomodulatory ability to their native equivalents.

### MPI and MRI bimodal detection of intra-myocardially injected magneto-iPSC-EVs

To evaluate the detectability of magneto-iPSC-EVs using MPI and MRI, we intra-myocardially injected 2 × 10^9^ magneto-iPSC-EVs (equivalent to 3.5 µg iron) into mouse heart subjected to ischemia-reperfusion injury (IRI). The experimental timeline is illustrated in **Figure 4a**, where in vivo MPI was performed at days 1 and 7 post-injection, followed by ex vivo MRI and histological analysis. Our results demonstrate that in vivo MPI provides sensitive detection of magneto-iPSC-EVs in the heart for up to 7 days post-injection (**Figure 4b**). MPI signals were quantified by comparing the signal intensity of the heart to fiducial markers containing a known SuperSPIO20 concentration (200 ng in 100 µL). At day 1, the signal-to-fiducial ratio was determined to be 5.6:1, corresponding to a retention of approximately 1.12 µg Fe of SuperSPIO20 (32% retention rate). By day 7, the signal-to-fiducial ratio decreased by approximately 50%, suggesting a significant clearance of magneto-EVs from the myocardium (**Figure 4c**).

**Figure 4.**
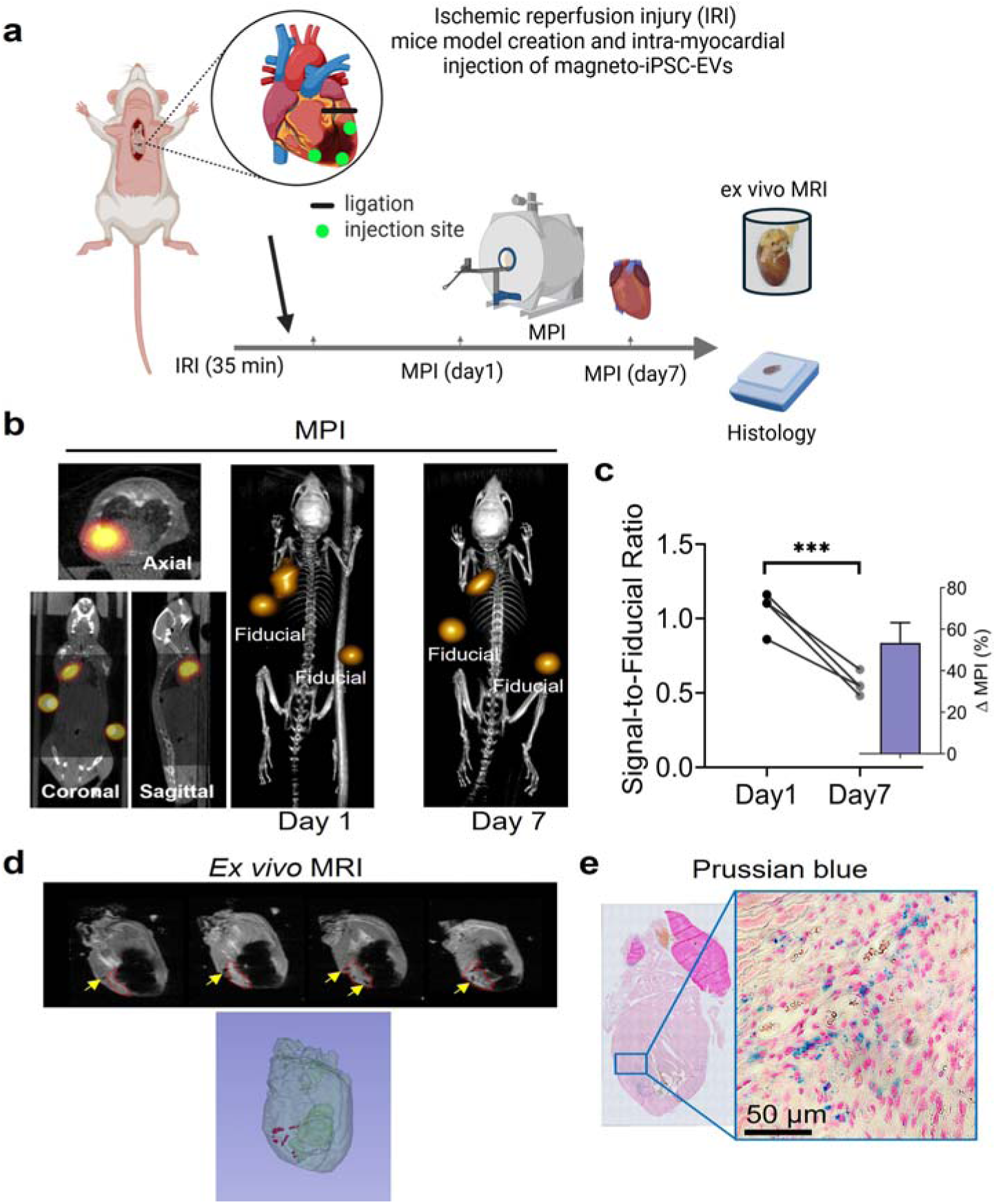
In vivo tracking of magneto-iPSC-EVs in a mouse model of myocardial ischemia-reperfusion injury (IRI) using MPI and MRI. (**a**) Schematic illustration of the experimental timeline, showing the model of IRI model induction, intramyocardial injection of magneto-iPSC-EVs, and subsequent imaging time points. The inset depicts the site of coronary artery ligation and injection. (**b**) Representative 2D MPI views at day 1(left), 3D MPI image at day 1 (middle), and 3D MPI image at day 7 (right), demonstrating the magneto-iPSC-EVs distribution, with fiducial markers (200 ng SuperSPIO20 in 100 µL PBS) for signal reference. (**c**) MPI signal intensity change (quantified by signal-to-fiducial ratio) at day 1 and day 7 in each mouse, demonstrating a significant decrease over time (***P < 0.001). Inset shows the averaged percentage change in MPI signal after 7 days (n=3). (**d**) Ex vivo MRI images of IRI mouse heart 7 days post-injection, with yellow arrows indicating the presence of magneto-iPSC-EVs. A 3D reconstruction of the heart is shown at the bottom, with red dots representing the distribution of magneto-iPSC-EVs. (**e**) Histological verification of magneto-iPSC-EV retention using Prussian blue staining. The left panel shows a low-magnification view of the heart section, while the right panel presents a high-magnification (50 µm scale bar) image demonstrating SPIO (blue) within myocardial tissue.

By the same magnetic labels, MRI can also be used to visualize magneto-EVs, offering a complementary method to MPI for high-resolution imaging. In this study, we demonstrated the ability of (ex vivo) MRI to evaluate the distribution of magneto-iPSC-EVs within the heart (**Figure 4d**). The results revealed that the remaining magneto-EVs were localized at the three injection sites near the infarcted regions. A 3D reconstruction further confirmed this distribution pattern. While only ex vivo MRI was performed in this study, there are no technical barriers to performing in vivo MRI in future experiments.

The findings from MPI and MRI studies were validated by Prussian blue staining, which confirmed the presence of iron-labeled EVs primarily in the apex of the heart (**Figure 4e**). Notably, sparse blue dots were observed outside the injection sites, suggesting potential uptake and redistribution by host macrophages or other immune cells. Together, these results highlight the utility of MPI for quantifying EV retention over time and MRI for visualizing their spatial distribution within cardiac tissue as two complementary imaging modalities.

### Therapeutic effects of magneto-iPSC-EVs

As illustrated in **Figure 5a**, the therapeutic potential of magneto-iPSC-EVs, along with their native counterparts, was evaluated using cardiac MRI and Masson’s trichrome staining 30 days after a single-dose treatment. Representative short-axis MR images from each group at the end-diastole and end-systole phases of the cardiac cycle are shown in **Figure 5b**. In saline-treated mice, myocardial infarction (MI) resulted in a significantly enlarged end-systolic left ventricular (LV) volume compared to the sham group, indicating pathologic ventricular remodeling often associated with impaired contractile function. Treatment with both iPSC-EVs and magneto-iPSC-EVs effectively reversed this LV enlargement, restoring volumes to a level comparable to the sham control. As shown in **Figure 5c**, magneto-iPSC-EVs significantly improved left ventricular ejection fraction (LVEF) by 37.3% compared to saline control group (60.2 ± 10.0 vs 43.9 ±10.6, P = 0.0014), while native iPSC-EVs showed slightly higher LVEF (64.8 ± 7.3), corresponding to 47.6% improvement (P < 0.0001). No significant difference was observed between the magneto-iPSC-EVs and native iPSC-EVs groups (P = 0.5117), suggesting comparable efficacy between the two treatments.

**Figure 5.**
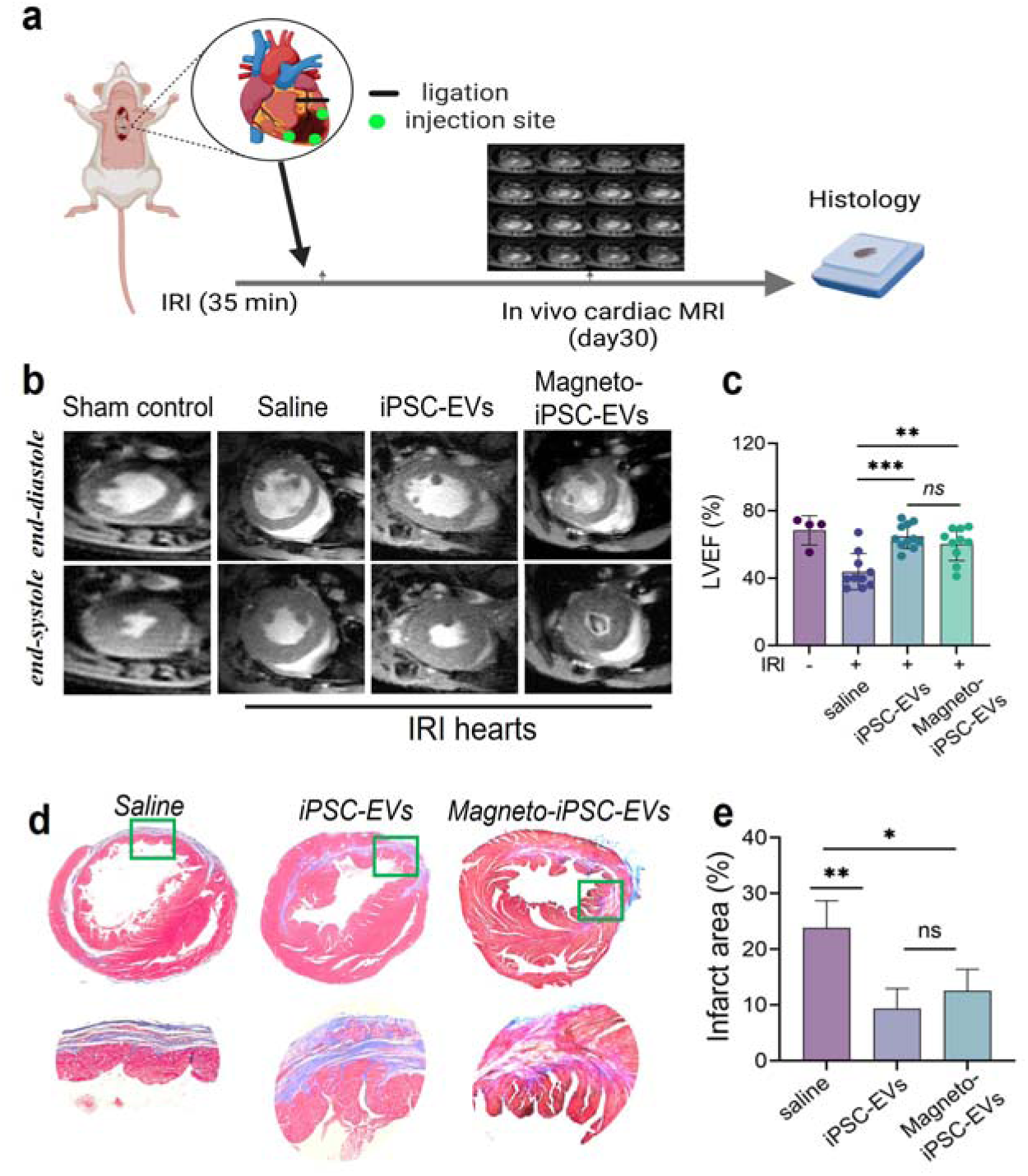
Therapeutic efficacy of magneto-iPSC-EVs in mouse hearts following IRI. (**a**) Experimental workflow showing IRI induction (35 min ischemia), intramyocardial delivery of therapeutic agents, and subsequent cardiac MRI and histological analysis at day 30. (**b**) Representative cardiac MR images showing mid-ventricular short-axis views during end-diastole and end-systole in sham control and IRI hearts treated with saline, iPSC-EVs, or magneto-iPSC-EVs. (**c**) Bar plots of left ventricular ejection fraction (LVEF%) in treatment groups, demonstrating significant improvement in EV-treated groups compared to saline controls (***P < 0.001, **P < 0.01, ns: not significant). (**d**) Masson’s trichrome staining of heart sections, with fibrotic tissue appearing blue and viable myocardium in red. Upper panel shows cross-sectional views; lower panel shows a zoomed-in view of the apical area indicated by the boxes. (**e**) Quantification of infarct size expressed as percentage of fibrotic area relative to total myocardial area (*P < 0.05, **P < 0.01, ***P < 0.001, ns: not significant).

To further evaluate therapeutic outcomes, Masson’s trichrome staining was used to assess fibrosis (blue) relative to viable myocardium (red). The results revealed a significant reduction in fibrosis area in both iPSC-EVs and magneto-iPSC-EVs groups compared to the saline group (**Figure 5d**). Higher magnification images highlighted the thin ventricular wall in the saline group, whereas both native- and magneto-iPSC-EV-treated mice exhibited thicker LV walls with more viable myocardium and less fibrotic tissue. As summarized in **Figure 5e**, quantitative analysis showed that the mean fibrosis area was 63.1 ± 14.1% in the saline group, 35.0 ± 8.7% in the iPSC-EVs group, and 32.2 ± 11.2% in the magneto-iPSC-EVs group. Compared to saline-treated mice, fibrosis was reduced by 44.6% (P = 0.0059) in the iPSC-EVs group and by 49.0% (P = 0.0182) in the magneto-iPSC-EVs group, with no significant difference between the two EV treatments. These findings demonstrate that magneto-iPSC-EVs exhibit therapeutic effects comparable to native iPSC-EVs, significantly enhancing viable myocardial mass and reducing infarct size.

## Discussion

In this study, we have established a novel theranostic platform by integrating MPI/MRI bimodal imaging with iPSC-derived EVs and demonstrated its application in treating myocardial infarction. We first optimized the electroporation protocol to achieve high SuperSPIO20 loading efficiency while minimizing the impact on EV integrity and therapeutic potency. Next, we conducted in vitro and in vivo studies to assess the MRI and MPI imaging capabilities of the prepared magneto-EVs. The imaging results clearly illustrated the excellent quantification provided by MPI, along with high-resolution images from MRI, making the two complementary techniques. Furthermore, compared to native iPSC-EVs, the therapeutic effects of magneto-iPSC-EVs are largely preserved, as evidenced by both in vitro assays and in vivo therapeutic evaluations. This new theranostic EV platform offers imaging tools for non-invasively measuring the tissue-specific uptake and targeted delivery in the disease site. We chose iPSC-derived EVs because of their reported therapeutic effects on ischemic myocardium^18,55^, and high EV production rate (16 folds higher than MSCs ^56,57^). The optimized magnetic labeling and imaging protocols can be easily tailored to other types of stem cell-derived EVs, including MSC-EVs.

In our previous study^39^ we developed a novel strategy to prepare highly purified, magnetically labeled EVs, termed magneto-EVs, which enable MRI tracking of the delivery of therapeutic EVs to the sites of injury. Our approach is simple, less equipment-intensive, and much faster (∼minutes) compared to other reported approaches^32,58^. In the present study, we continued optimizing the protocols to achieve even higher labeling efficiency while preserving the functionalities of the native EVs. Our optimized electroporation protocol allows efficient labeling of iPSC-EVs at a loading efficiency of 1.7 ng Fe per 10^6^ EV, which allows sensitive detection of as low as 10^7^ EVs using either MPI or MRI, which is below the therapeutic threshold concentration levels. Compared to previous studies, the labeling efficiency is significantly higher. For instance, in a recent study by Toomajian, approximately 0.5 fg Fe per EV (corresponding to 0.5 ng Fe per 10^6^ EVs) was obtained in 4T1 tumor cell-derived EVs using an indirect labeling approach^33^. This essentially is 7 times lower than our loading efficiency. While different SPIO particles were used, making a direct comparison difficult, our approach offers almost one order of magnitude of higher sensitivity for EV detection. Another advantage of our approach is that the prepared magneto-EVs are in high purity because unencapsulated SuperSPIO20 nanoparticles can be removed efficiently by Ni-NTA thanks to the his-tags conjugated to the surface of SuperSPIO20. The high purity is crucial for passing quality control requirements in future translational studies.

It should be noted that not only electroporation pulses but also electroporation buffer can strongly affect the quality of the resulting EVs. In our study, we selected KPBS over the commonly used PBS because research has shown that EVs stored or processed in PBS are prone to aggregation, fusion, or morphological changes, whereas those in KPBS demonstrate improved preservation of EV size, shape, and bioactivity^59^. Furthermore, natural, non-toxic disaccharides such as sucrose and trehalose could preserve the stability and morphology of EVs while minimizing aggregation as membrane stabilizers, particularly under stress conditions like electroporation. Our results show that 50 mM trehalose in KPBS outperforms other combinations, providing the best protection for EVs undergoing electroporation.

Our results show the great potential of magnetic labeling for enabling non-invasive imaging of magneto-iPSC-EVs in the infarcted mouse heart. MPI provided hot-spot images with zero background signal, enabling highly sensitive detection and quantification of the injected EVs. Based on the MPI signal ratio between the heart and fiducial markers, we estimated that approximately 32% of the injected EVs were retained in the heart at 24 hours post-injection, and 50% of those (16% of the total amount of EVs injected) remained in the heart by day 7. High-resolution ex vivo MRI further revealed the focal distribution of magneto-iPSC-EVs at the injection sites, which was validated by Prussian blue staining. These imaging results demonstrate the capability of this bimodal imaging approach to non-invasively assess the biodistribution and retention of therapeutic EVs, which is critical for guiding the development and optimization of EV-based therapies.

Importantly, magneto-iPSC-EVs exhibited significant therapeutic efficacy in a mouse model of myocardial infarction. Cardiac MRI evaluation showed that magneto-iPSC-EV treatment significantly improved LVEF and reduced scar size compared to the saline control group. Histological assessment further revealed increased viable myocardium and reduced fibrosis in the magneto-iPSC-EV treated group. These results are consistent with previous studies reporting the cardioprotective and regenerative effects of iPSC-EVs in animal models of myocardial infarction^14,18^. The preservation of therapeutic efficacy in magneto-iPSC-EVs highlights the potential of this theranostic approach to enhance the development and translation of EV-based therapies without compromising their inherent therapeutic properties. Compared to direct injection of parental iPSCs or MSCs, magneto-iPSC-EVs offer several advantages, including reduced risks of tumorigenicity and immune rejection associated with cell-based therapies. Additionally, while cell therapies often face challenges related to poor engraftment and survival in the ischemic myocardium, EVs can exert their therapeutic effects through paracrine signaling without requiring long-term retention in the target tissue, potentially leading to more consistent outcomes.

The choice of the SPIO nanoparticle type is a critical factor in optimizing the performance of magneto-EVs for MPI. In this study, we used dendronized SuperSPIO20 nanoparticles with a core diameter of 20 nm, which is considered an optimal size for MPI applications. Tay et al. demonstrated that SPIOs with iron oxide core diameters between 15-25 nm exhibit the highest MPI signal intensity and resolution^60^. The slight increase in the full-width at half-maximum (FWHM) of the MPI signal for magneto-iPSC-EVs (50 mT) compared to unencapsulated SuperSPIOs (20 mT) can be attributed to the increased Brownian relaxation of EV-encapsulated SPIOs, as the matrix composition of the surrounding medium can affect both the MPI signal amplitude and FWHM value^61^. Despite this spectrum-broadening effect, the MPI signal remained strong and linearly correlated with the concentration of magneto-iPSC-EVs, enabling quantitative tracking of the administered EVs in vivo.

In this study, we utilized intramyocardial injection for delivering magneto-iPSC-EVs to the injured myocardium. This approach offers numerous advantages in the context of treating myocardial infarction^62^. Firstly, direct intramyocardial injection allows for precise targeting of the peri-infarct areas, ensuring a high local concentration of therapeutic EVs at the site of injury. This targeted delivery is particularly crucial in the acute phase of myocardial infarction when rapid intervention is necessary to minimize tissue damage and promote repair. The operation can be guided by interventional X-ray, ultrasonic imaging, and MRI^63^. In our study, we used MPI/MRI to reveal a substantial retention rate of 32% after 24 hours, with detectable signals persisting for up to 7 days, when an intramyocardial route was used. The sustained presence of EVs in the infarcted area is essential for their therapeutic efficacy. However, it is important to acknowledge that intramyocardial injection is an invasive procedure, which may limit its clinical applicability in certain scenarios. Future studies should explore less invasive delivery routes such as intracoronary or intravenous administration, potentially combined with targeting strategies to enhance myocardial uptake. Comparing the biodistribution, retention, and therapeutic efficacy of magneto-iPSC-EVs delivered via different routes will provide the basis for optimizing this theranostic approach for clinical translation.

While our study demonstrates the potential of magneto-iPSC-EVs as a theranostic platform for MI, several limitations should be acknowledged. Firstly, the magnetic labeling process resulted in a significant increase in EV size, which may potentially impact the pharmacokinetics and biodistribution of the EVs in vivo when a systemic injection is used. Future studies should investigate whether the altered size affects the EVs’ ability to cross biological barriers or their cellular uptake mechanisms. Moreover, our current imaging approach relies on detecting the SPIO nanoparticles rather than the EVs themselves. This presents a limitation for long-term monitoring, as the SPIO could potentially be released from the EVs in vivo over time. Addressing these limitations in future research will be crucial for advancing the clinical translation of magneto-iPSC-EVs as a theranostic tool for myocardial infarction and other ischemic diseases.

## Conclusions

In this study, we have developed and refined a magnetic labeling protocol for producing MPI/MRI-trackable iPSC-EVs and evaluated it in a mouse MI model. The distribution, quantity, and retention of injected EVs were visualized using non-invasive MPI and MRI. Despite the increased EV size and cargo loss resulting from magnetic labeling procedures, the intrinsic regenerative potential was largely preserved. This novel theranostic EV platform may guide future development and clinical translation of EV-based therapy for treating ischemic cardiovascular disease.

## Acknowledgments

This work was supported by NIH R33 HL161756, R01 CA261974, S10 OD026740, P41 EB024495, and 2023-MSCRFL-6121. W Wang is a recipient of the MSCRF2024 fellowship (2024-MSCRFF-6377).

## Data Availability Statement

Data presented in this study are available on request from the corresponding author.

