## Supporting information for "Magnetically Labeled iPSC-Derived Extracellular Vesicles Enable MRI/MPI-Guided Regenerative Therapy for Myocardial Infarction"

iPSCs were thawed and passaged onto vitronectin-coated 6-wells plates (250,000 cells/well) in complete Essential 8™ medium (E8, Gibco, 2 mL/well) supplemented with 10 µM of Y-27632 ROCK inhibitor (Ri, STEMCELL Technologies), as previously described[1,2]. The E8 medium without Ri was replaced daily. Cells were passaged every 3 days upon reaching ~80-90% confluency (**Figure S1**). They were cultured for 10-12 passages.

Cultured cells were monitored daily for morphological characteristics to ensure the maintenance of healthy, undifferentiated colonies. Regular quality control assessments were performed to ensure the integrity and pluripotency of the iPSC line. Genomic stability was evaluated using G-band karyotyping and single nucleotide polymorphism (SNP) arrays. Pluripotency markers Oct4 and Tra-1-60 were assessed by immunohistochemistry (IC). Functional pluripotency was validated through trilineage differentiation assays. Cells were routinely checked for mycoplasma contamination.

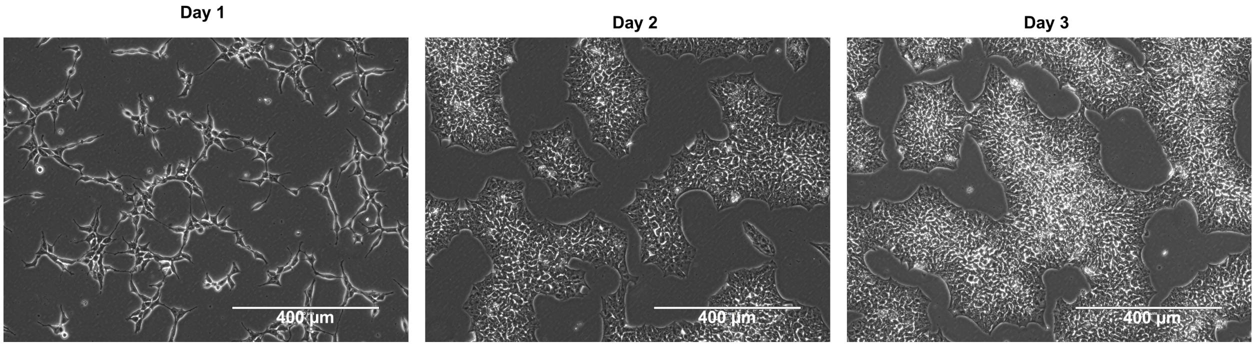


**Figure S1**. **Representative brightfield images of the BC1-iPSCs in culture.**

***S2. Western blots for EV characterization***

iPSC-EVs and magneto-iPSC-EVs samples (18 µL) were lysed in 1x radioimmunoprecipitation assay buffer (RIPA, Cell Signaling Technology,Cat. #9806) for overnight (approx. 18 hours) at 4°C. Lysates were heated at 95 °C for 5 minutes together with 6.67 µL 4x Laemmli Sample Buffer (Bio-Rad, #1610747; non-reducing condition). Lysates were resolved using a 4% -15% Criterion TGX Stain-Free Precast gel (Bio-Rad, # 5678084), with Spectra Multicolor Broad Range protein ladder (Thermo Scientific, # 26634). Stain-free images of a gel were obtained using Bio-Rad Gel Doc imager. Proteins were then transferred onto a PVDF membrane (Invitrogen, # IB24001) using iBlot 2 semi-dry transfer system (Invitrogen). Blots were first probed using primary antibodies, including CD9 (1:1000, Ms, BioLegend #312102), Calnexin (1:1000, Rb, Abcam #22595), in PBST (PBS with 0.05% Tween-20 (BioXtra, #P7949) and 5% Blotting Grade Blocker (Bio-Rad, #1706404). Then, SuperSignal West Pico PLUS Chemiluminescent Substrate (Thermo Scientific, # 34580) was applied to the membrane, and blots were imaged using an iBright 1500FL Imager (Thermo Fisher). A representative Western Blot image is provided in **Figure S2**, showing the presence of EV-positive marker CD9 and absence of EV-negative marker Calnexin.

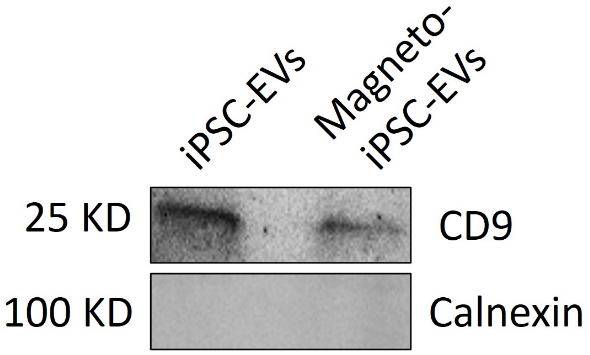


**Figure S2. Representative western blot of iPSC-EVs and magneto- iPSC-EVs.**

### *S3. Quantitative RT-PCR*

Primer sequences used in qRT-PCR are listed in **Table S1**.

**Table S1. Primer sequences used for RT-PCR**

| Gene | Primers | |
|---|---|---|
| | Forward | Reverse |
| TGF-β [3] | GGCCAGATCCTGTCCAAGC | GTGGGTTTCCACCATTAGCAC |
| Arg-1 [4] | GACCGTTGTGTGTGTTCTGG | GATGAGCAGCATCACAAGGA |
| IL–6 [5] | CTGCAAGAGACTTCCATCCAGTT | AGGGAAGGCCGTGGTTGT |
| 18S [6] | GTAACCCGTTGAACCCCATT | CCATCCAATCGGTAGTAGCG |
| Mcp1 [7] | CCACTCACCTGCTGCTACTCA | TGGTGATCCTCTTGTAGCTCTCC |

### *S4. Characterization of EVs after electroporation*

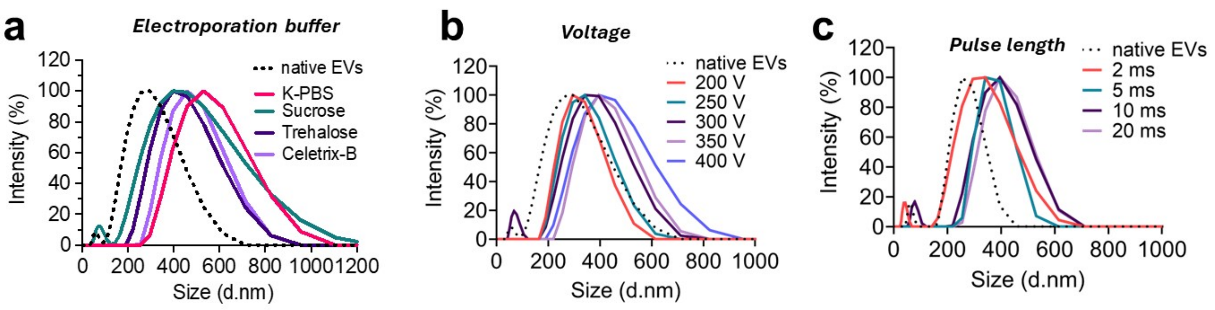


**Figure S3.** Effect of (**a**) buffer (pulse fixed at 400 V/10 ms), (**b**) pulse voltage (fixed pulse duration =10 ms), and (**c**) pulse length (fixed pulse voltage = 300 V) on the size of iPSC-EVs after electroporation as revealed by DLS measurements. The result shows that electroporation conducted in trehalose-containing KPBS buffer resulted in the least size increase (435.0±6.1 nm), followed by Celetrix buffer (477.0±24.9 nm), sucrose-containing KPBS (488.9±73.8 nm), and KPBS (584.9±61.1 nm), as compared to native EVs (293.4±29.9 nm).

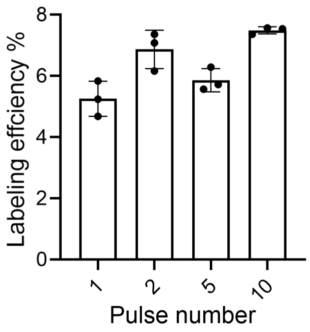


**Figure S4. Effect of pulse number on magnetic labeling efficiency in iPSC-EVs.** Electroporation pulses were applied at 300 V with the total duration kept at 10 ms.

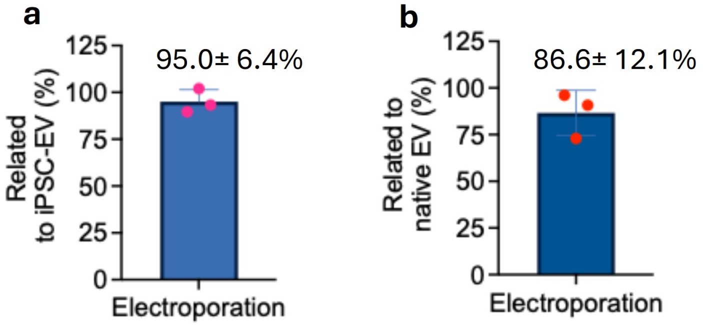

**Figure S5**. Characterization of cargo loss (percentage) of (**a**) Protein and (**b**) RNA contents in iPSC-EVs caused by the optimized electroporation (300 V, 10 ms, and 2 pulses).

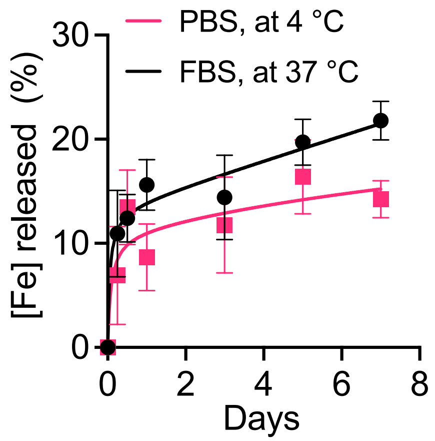


**Figure S6. Stability of magnetic labeling as characterized by the** released [Fe] from magneto-iPSC-EVs after incubating PBS (room temperature) or serum (FBS, 37 °C) at a volume ratio of 1: 2 for up to 7 days.

### *S5. Small RNA-sequencing analysis*

The top 20 abundant miRNAs in iPSC-EVs/magneto-iPSC-EVs identified by small RNA-seq are summarized in **Table S2**, along with their functions.

**Table S2. Summary of highly expressed miRNA in iPSC-EVs/magneto-iPSC-EVs.**

| ***Order of abundance*** | **Gene name** | **Function (reference)** |
|---|---|---|
| *1* | hsa-miR-148a-3p | Anti-inflammation [8] |
| *2* | hsa-miR-21-5p | Anti-apoptosis and anti-inflammation [9,10] |
| *3* | hsa-miR-92a-3p | Anti-apoptosis [11] |
| *4* | hsa-miR-151a-3p | Anti-apoptosis [12] |
| *6* | hsa-let-7f-5p | Anti-apoptosis and anti-inflammation [13,14] |
| *8* | hsa-miR-34c-5p | Anti-apoptosis and anti-inflammation [15] |
| *9* | hsa-let-7i-5p | Anti-apoptosis and anti-inflammation [16-18] |

| | | |
|---|---|---|
| *11* | hsa-let-7g-5p | Anti-apoptosis and anti-inflammation [19,20] |
| *13* | hsa-miR-30d-5p | Anti-apoptosis and anti-inflammation* [21] |
| *14* | hsa-miR-143-3p | Anti-apoptosis [22] |
| *15* | hsa-miR-26a-5p | Anti-apoptosis and anti-inflammation* [23] |
| *16* | hsa-let-7a-5p | Anti-fibrotic and anti-inflammation [24] |
| *17* | hsa-miR-423-3p | Anti-apoptosis* [25] |
| *19* | hsa-miR-27b-3p | Anti-apoptosis and anti-inflammation [26] |
| *20* | hsa-miR-3184-3p | Anti-apoptosis and anti-inflammation [27] |

* Previously demonstrated for myocardial infarction treatment.

### *S6. Ischemia-reperfusion injury (IRI) mouse model*

The cardiac ischemia-reperfusion injury (IRI) model was performed according to previously published protocols[28]. In brief, mice were anesthetized using 3%–4% isoflurane for induction and 0.03–0.07 mg/kg buprenorphine (subcutaneous) for pre-operative analgesia. Anesthesia was maintained with 1%–2% isoflurane and 2 mg/kg succinylcholine (intraperitoneal). The mice were intubated, mechanically ventilated, and maintained at a constant body temperature throughout the procedure.

A left thoracotomy was performed to expose the heart, and the left anterior descending (LAD) coronary artery was occluded for 35 minutes using a 7-0 PROLENE suture and PE10 tubing. The chest cavity was temporarily closed during the occlusion period. Five minutes before the end of occlusion, the chest was reopened, and the suture was removed to allow reperfusion. After closure of the chest, a second dose of buprenorphine (0.06–0.075 mg/kg, subcutaneous) was administered for post-operative analgesia. Sham-operated mice underwent the same surgical procedures without LAD ligation.
